# In vivo AGO-APP identifies a module of microRNAs cooperatively controlling exit from neural stem cell state

**DOI:** 10.1101/2023.10.05.560992

**Authors:** Karine Narbonne-Reveau, Andrea Erni, Norbert Eichner, Surbhi Surbhi, Gunter Meister, Christophe Beclin, Cédric Maurange, Harold Cremer

## Abstract

MicroRNAs (miRNAs) are essential regulators of all developmental processes. Their function is particularly important during neurogenesis, when the production of large numbers of neurons from a limited number of neural stem cells depends on the precise control of determination, proliferation and differentiation. However, miRNA regulation of target mRNAs is highly promiscuous, one miRNA can target many mRNAs and vice versa, raising the question of how specificity is achieved to elicit a precise regulatory response.

Here we introduce AGO-APP, a novel approach to purify Argonaute-bound miRNAs directly from cells and tissues in vivo, to isolate actively inhibiting miRNAs from different neural cell populations in the larval Drosophila central nervous system. We identify a defined group of miRNAs that redundantly target all iconic genes known to control the transition from neuroblasts to neurons. In vivo functional studies demonstrate that knockdown of individual miRNAs does not induce detectable cellular phenotypes. However, simultaneous knockdown of multiple miRNAs leads to precocious stem cell differentiation, demonstrating functional interdependence. Thus, miRNAs cooperate within a regulatory module to specify the targeted gene network.

## Introduction

MicroRNAs (miRNA) suppress protein production post-transcriptionally by binding to a short sequence motif in the 3’UTR of target mRNAs. Bioinformatic tools predict that a single miRNA can target hundreds of mRNAs and, vice versa, a single mRNA can be regulated by several miRNAs^1–3^, indicating that miRNA/mRNA interactions are intrinsically promiscuous. Moreover, loss of specific miRNAs induces generally only minimal changes in target gene expression^4,5^ and often only mild to absent phenotypes ^6,7^.

In light of these observations, it is largely unclear how target specificity and silencing efficiency is achieved in a given cell type to induce a particular biological function. It has been hypothesized that additive, synergistic, or even combinatorial interactions contribute to this process^8,9^. However, this aspect is particularly hard to experimentally address, requiring detailed insight into the expression of functional miRNAs and their targets, as well as the development of new technologies and approaches for functional analyses.

Brain development is a highly orchestrated process involving the generation of thousands of different types of neurons and glial cells from a relatively small number of neural stem cells (NSCs). Their coordinated production and maturation ultimately lead to the formation of highly complex functional networks. Not surprisingly, the mammalian brain represents the organ with the highest miRNA expression levels and complexity ^10,11^, and various specific miRNAs have been functionally implicated in basically all processes leading from NSCs to functional neurons^12–14^.

Additive or cooperative effects are hard to study in vertebrates due to the complex and time-consuming nature of genetic manipulations. *Drosophila* represents a comparably simple and accessible model for neurobiological research^15^. NSCs and their lineage are described in unmatched detail and powerful genetic tools allow to manipulate with exquisite precision the expression of any gene or miRNA. *Drosophila* NSCs, known as neuroblasts, divide asymmetrically throughout developmental stages to self-renew while generating intermediate progenitors, called Ganglion Mother Cells (GMCs)^16^. GMCs usually undergo a single division to generate two neurons or glia. This process involves the activity of key cell fate determinants including the transcription factors Prospero and Nerfin-1, as well as the RNA-binding proteins Brat and Elav. GMCs lacking Prospero, Nerfin-1 or Brat fail to differentiate and reacquire a neuroblast identity, leading to the generation of ectopic neuroblasts^17–20^. Elav, an RNA binding protein, is expressed as soon as post-mitotic neurons are produced and is a general marker for neuronal differentiation^21^. In addition, as neuroblasts undergo successive rounds of asymmetric divisions during embryonic and larval stages, they transit through temporal windows that regulate self-renewing potential and the identity of neurons and glia as they are being produced^22,23^. Genes defining temporal windows include the transcription factors Chinmo, Mamo and Eip93F, as well as the RNA-binding proteins Imp and Syncrip^24^. Their deregulation affects neuronal and glial identity as well as neuroblast self-renewal. Thus, the highly coordinated control of asymmetric division and temporal patterning, based on a group of well-defined factors, ensures that each neuroblast generates its correct repertoire of neuronal and glial progeny by the end of development. It appears likely that, like in vertebrates, miRNA regulation represents a superposed regulatory level in this process. However, in the fly nervous system only few candidate miRNAs have been studied in detail and implicated in the control neuronal/glial identity and function and for regulating neuroblast self-renewal^14^. A comprehensive view of miRNA expression during *Drosophila* nervous system development is lacking, precluding a detailed and integrated understanding of miRNA function and mode of action during neurogenesis.

Here, we adapt the Ago protein Affinity Purification by Peptides (AGO-APP) technology for its in vivo use. This approach is based on the transgenic expression of a small peptide of about 80 AA, derived from RISC complex protein GW182, that binds to all known AGO proteins with high affinity, allowing their immunoaffinity mediated pulldown in complex with the bound miRNAs that can be subsequently released and analyzed^25^. Using this approach, we provide a comprehensive list of miRNAs purified from neuroblasts, neurons and glial cells of the larval *Drosophila* central nervous system (CNS). Bioinformatic analyses identified a regulatory module of miRNAs targeting all key genes that control neuroblast-to-neuron transition as well as most temporal patterning genes. To investigate the function and mode of action of this miRNA/mRNA module, we compared the consequences of inactivation of individual miRNAs members, to simultaneous knockdown of groups of miRNAs, demonstrating that miRNAs cooperate to regulate NB self-renewing potential versus differentiation. Our study suggests that the concerted action of several miRNAs allows the silencing of a defined ensemble of genes involved in a specific biological process.

## Results

### Ago-APP identifies cell type specific miRNAs in larval neurogenesis

T6B is a small peptide of 85 AA that is derived from the GW182 family protein TNRC6B, an active component of the RISC complex. T6B binds with high affinity to all known Argonaute (AGO) proteins^25^. Based on this strong binding we developed AGO-APP. This approach relies on the expression of the T6B peptide tagged with FLAG-HA-YFP (T6B-FHY) (Fig. 1A). Binding of T6B to AGO disrupts the RISC by displacing GW182 proteins. Subsequently, the remaining miRNA:Ago:T6B-FHY complex can be pulled down with anti GFP-nanobodies, the miRNA is released and can be analyzed by RT-qPCR or sequencing^25^ (Fig. 1A).

**Figure 1:**
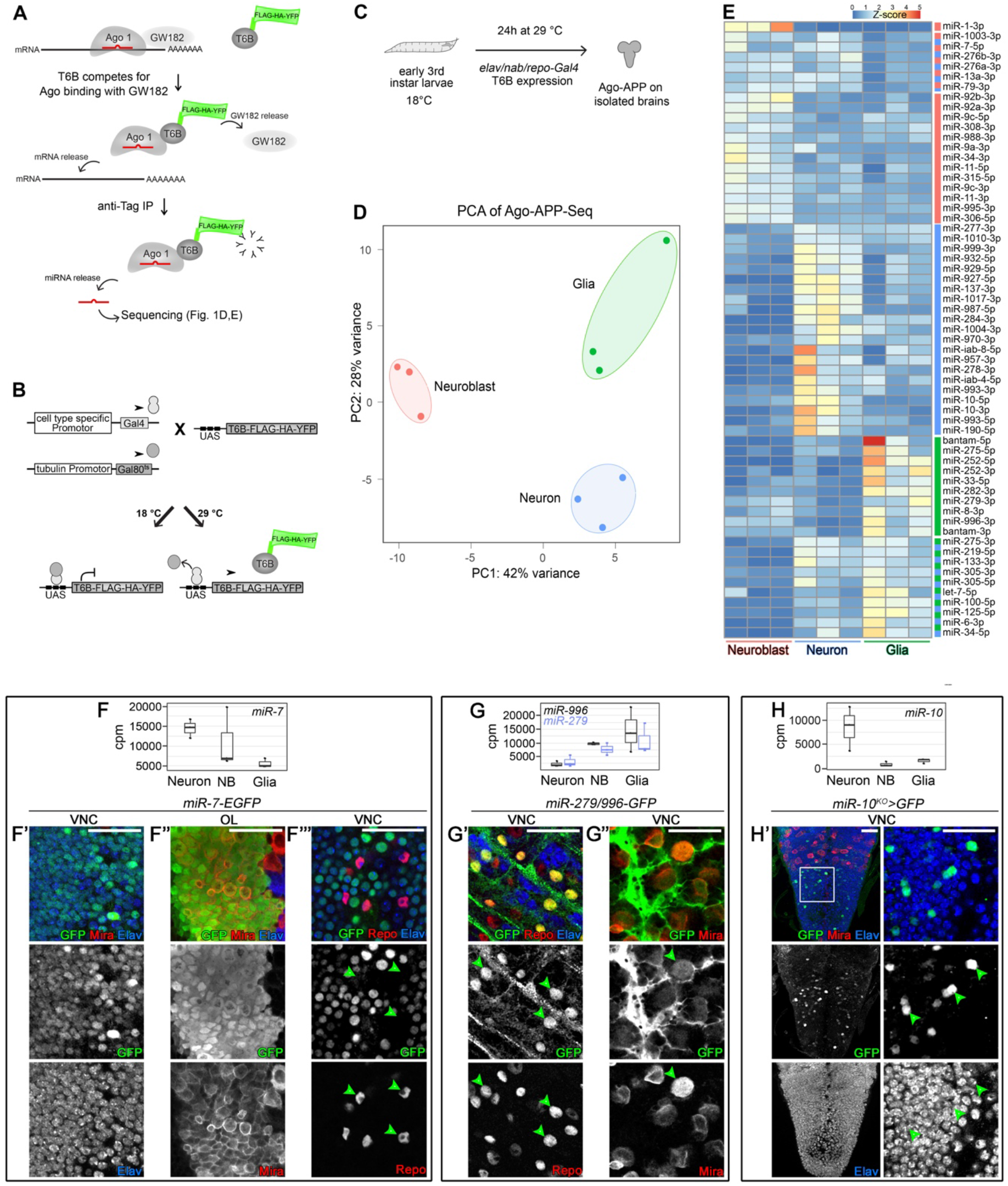
Ago-APP isolates cell-type specific miRNAs. **A)** The Ago-APP procedure. **B)** Spatio-temporal control of *T6B-FLAG-YFP-HA (T6B-FYH*) transgene expression using the UAS/GAL4/GAL80ts system in *Drosophila*. **C)** Schematic representation of cell type-specific Ago-APP in *Drosophila* larvae. **D)** PCA analysis of sequencing data after Ago-APP was performed on neuroblasts, neurons or glia (three independent replicates per condition). **E)** Heat-map of normalized values (computed by dividing the normalized read counts by the mean of all samples) for each miRNA showing a DeSeq2 differential expression between 2 conditions. **F)** Box plots of normalized miR-7-5p read counts in Ago-APP samples from neurons, neuroblasts (NB) and glial cells. **F’-F’’)** Immunostaining against GFP, Mira and Elav in the VNC (F’) or in the OL (F’’) of a *miR-7-EGFP* reporter *Drosophila* line. F’”) Immunostaining against GFP, Repo and ELAV in the VNC of a *miR-7-EGFP* reporter *Drosophila* line. Arrowheads show cells positive for Repo but negative for GFP. **G)** Box plots of normalized miR-279-3p and miR-996-3p read counts Ago-APP samples from neurons, NBs and glial cells. **G’-G’’)** Immunostaining against GFP, Repo and Elav (G’) or Mira (G’’) of a miR-279/996*-GFP* reporter *Drosophila* line. **H)** Box plots of normalized miR-10-3p read counts in Ago-APP samples from neurons, NBs and glial cells. **H’)** Immmunostaining against GFP, Mira and Elav in the VNC of a *miR-10-GAL4; UAS-nuclearEGFP* reporter *Drosophila* line. Only a subset of Elav positive neurons is positive for GFP (shown with arrowhead). The scale bars represent 30 µm.

We aimed at using T6B *in vivo* to isolate Ago bound miRNAs directly from specific cell types of *Drosophila melanogaster*. A transgenic *Drosophila* line expressing T6B-FHY under the control of *UAS* sequences was generated, allowing cell-type specific expression of the fusion protein when combined with suited *GAL4* driver lines (Fig.1B)^26^. For temporal control, we combined this approach with the *GAL80* temperature sensitive system (*tubulin-GAL80^ts^*)^27^. Thus, when *x-GAL4, tub-GAL80^ts^*, *UAS-T6B* animals are maintained at 18° (restrictive temperature), *GAL4*-dependent expression is prevented. Switching to 29° (permissive temperature) for 24 hours allows *GAL4-*dependent expression (Fig.1B,C). For cell type specific expression, we combined this temporal control system with well characterized driver lines for i. neuroblasts (*nab-GAL4/tub-Gal80^ts^*)^28^, ii. the entire neuronal lineage comprising predominantly neurons (*elav-GAL4/tub-Gal80^ts^)*^29^ and iii. glial cells (*repo-GAL4/tub-Gal80^ts^*)^30^ (Fig.S1). For miRNA isolation, 20 late larval CNS were dissected 24h after induction of T6B expression for each transgenic combination. Following T6B-FHY pulldown, AGO-bound miRNAs were released and analyzed by high-throughput small RNA sequencing using a bias minimized sequencing protocol^31^. All experiments were performed in three biological replicates.

Principal component analysis revealed strong clustering of replicates for each cell type, demonstrating the high degree of reproducibility of the approach. Clear separation of all three clusters, indicated the expression of specific miRNA subsets in neuroblasts, neurons and glia (Fig.1D). Heat map presentation of all miRNAs that showed statistically significant differential expression between two samples in DEseq2 analyses^32^ identified neuroblast-, neuron- and glial cell-specific miRNAs subsets (Fig.1E).

Next, we validated the expression of selected cell-type enriched miRNAs, as identified by AGO-APP, using a set of reporter lines. AGO-APP showed that *miR-7* expression was strongest in neurons, present at lower levels in neuroblasts and quasi-absent in glia (Fig. 1F). In agreement, a *(miR-7)E>GFP* reporter^33^ showed high fluorescence levels in Elav-positive neurons of the ventral nerve cord (VNC) (Fig.1F’,F’”), while Repo-positive glial cells were always negative (Fig.1F’”, green arrowheads). Mira-positive neuroblasts of the optic lobe (OL) showed also GFP expression, as expected^34^ (Fig.1F”).

Mir-279 and miR-996, two miRNAs with related seed sequences and generated from a joint transcriptional unit, showed an opposing expression to miR-7 in our AGO-APP samples, being strong in isolates from glial cells, lower but substantial in the neuroblast fraction and barely detectable in neurons (Fig.1G). In agreement, a *miR-279/996-GFP* reporter^35^ showed highest fluorescence levels in Repo positive glial cells (Fig.1G’, arrowheads), moderate levels in Mira expressing neuroblasts (Fig.1G’’, arrowhead) and absence of label in Elav-positive neurons (Fig.1G’).

Finally, miR-10 was strongly present in sequenced neuronal isolates but was very low in neuroblasts and glia (Fig.1H). In agreement, a *miR-10^KO^-GAL4* line^36^ showed expression in a subpopulation of Elav-positive neurons in the ventral nerve cord (Fig.1H’, green arrowheads) but not in Mira-positive neuroblasts (Fig.1H’, red) or glial cells (not shown). This suggests that Ago-APP allows to detect miRNAs present in small cellular subpopulations in a large background of negative cells, pointing to the high sensitivity of the approach.

Altogether, these analyses demonstrate that AGO-APP represents a sensitive and reliable method that allows isolation of all AGO-bound miRNAs in a single step. Importantly, AGO-APP represents a benchtop approach that does not depend on cell isolation.

### A regulatory module controlling neuroblast-to-neuron transition

Next, we addressed the functional significance of the expression data. In mammals and *Drosophila*, miRNAs have been implicated in the control of stem cell maintenance versus differentiation^37–39^. Therefore, we focused on miRNAs that were downregulated during the transition from neuroblasts- to-neurons following asymmetric division (Fig. 2A). Sixteen miRNAs showed a significant (adjusted p-value < 0.05) downregulation in neurons compared to neuroblasts (Fig. 2B). Three pairs of these miRNAs (miR-279-3p/miR-996-3p; miR-11-3p/miR-308-3p, miR-92a-3p/miR-92b-3p) shared seed sequences and were grouped in further analyses. We used the Targetscan algorithm^1^ to determine the predicted target genes of each neuroblast-enriched miRNA. Target prediction, in combination with transcriptomic analysis previously performed in neuroblasts and neurons^40^, demonstrated that these neuroblast-enriched miRNAs predominantly target mRNAs encoding neuronal proteins (Fig. 2C), consistent with a differentiation inhibiting function.

**Figure 2:**
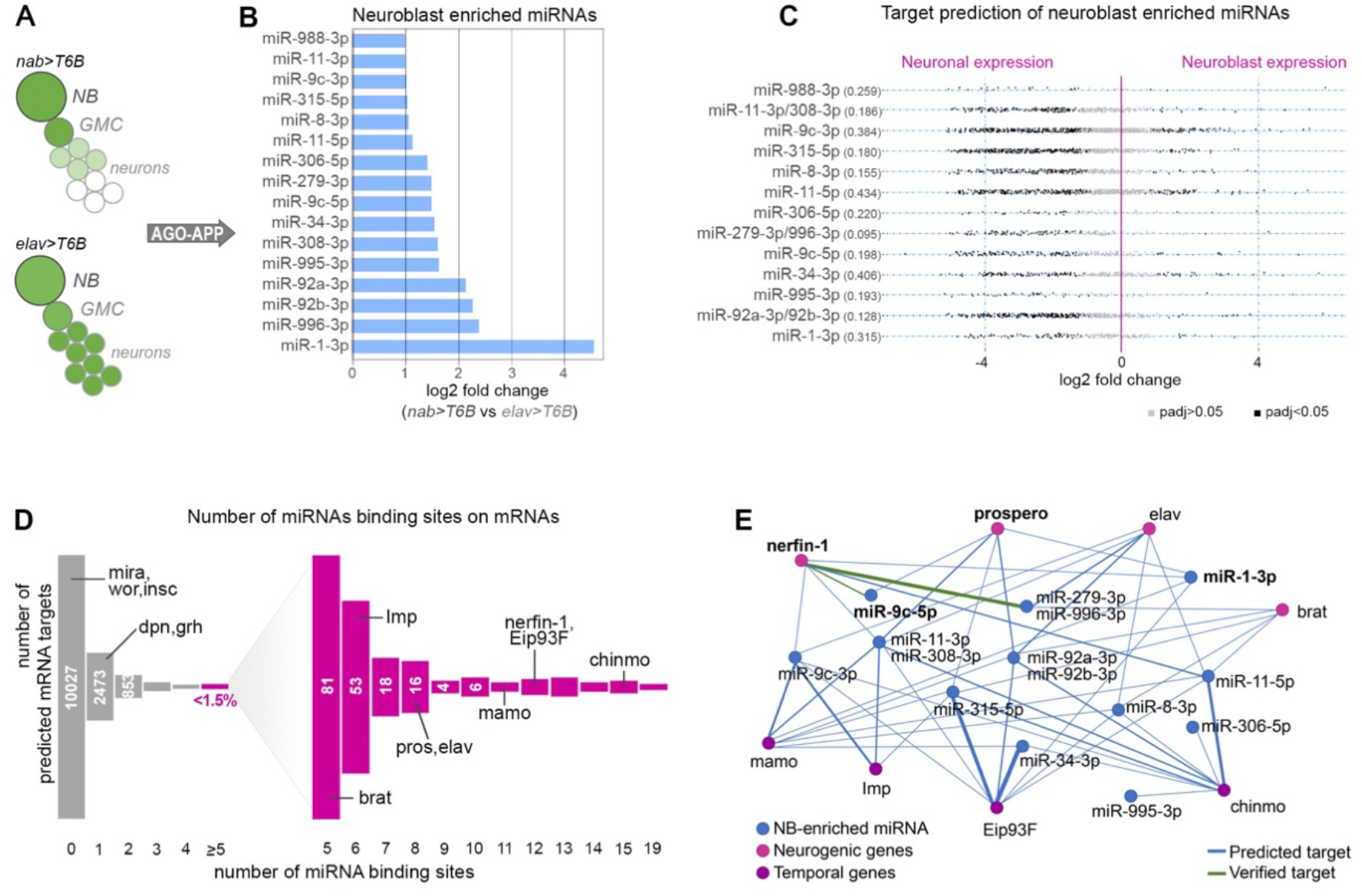
A neuroblast-enriched miRNA module targeting differentiation and temporal genes. **A)** Scheme of T6B expression in a neuroblast lineage when driven by *nab-GAL4* or *elav-GAL4*. **B)** Histogram showing the fold change of miRNAs that are significantly over-expressed in Ago-APP neuroblast samples compared to Ago-APP neuron samples. **C)** Scatter plot representing for each neuroblast-enriched miRNA the differential expression of each predicted target gene when comparing neuroblasts vs neurons, based on gene expression sequencing results published in Berger et al (2012) (see mat and methods). Dark grey dots correspond to target genes for which the differential expression is significant. In brackets is shown for each miRNA the ratio (number of neuroblast-enriched targets) / (number of neuron-enriched targets) **D)** Distribution of *Drosophila* genes along an axis representing the number of sites in their 3’UTR predicted to be targeted by the module of neuroblast-enriched miRNAs. Master genes involved in neuroblast maintenance (dpn, grh, mira, nab, wor, insc) are poorly or not targeted, while master neuronal differentiation (brat, pros, elav, nerfin-1) or temporal patterning genes (Imp, mamo, Eip93F, chinmo) are predicted to be highly targeted. **E)** Predicted interactome between the neuroblast-enriched miRNAs and neurogenic or temporal mRNAs. The blue lines correspond to predicted interactions according to the TargetScan algorithm, while green lines represent previously experimentally validated interactions. The thickness of the connecting lines is proportional to the number of times the mRNA is targeted by the miRNA.

MiRNA mediated regulation shows a high degree of promiscuity and non-functional interactions, rendering the identification of functional interactions difficult. In agreement, a genome-wide analysis using TargetScan predicted the large number of 4026 out of 14053 coding genes to be targeted at least once by one of the 16 neuroblast-enriched miRNAs (Fig. 2D). However, previous work provided strong evidence that the efficiency of regulation by miRNAs depends on the number of binding sites present in the 3’UTR^8,9^. Setting a limit of at least 5 target sites reduced the list of potential mRNAs to 201, representing less than 1,5% of all coding genes. Gene ontology analyses indicated that these genes are strongly enriched in biological processes involving development, neurogenesis and neuronal differentiation (Fig.S2A). Strikingly, the group of genes targeted ≥ 5 times by neuroblast-enriched miRNAs contained all iconic genes known to induce neuron differentiation after neuroblast asymmetric division^20^, including nerfin-1 (12 target sites), prospero (8 target sites), elav (8 target sites) and brat (5 target sites). In addition, well-described temporal patterning genes in larval neuroblasts^22,24,41^ were also repeatedly targeted by the same module of miRNAs, including Imp (6 target sites), mamo (11 target sites), Eip93F (12 target sites) and chinmo (15 target sites), most of them being silenced in late larval neuroblasts (Fig. 2D,E; FigS2B). In contrast, typical neuroblast-specific genes^16^, like miranda, worniu and inscutable, exhibited no target sites, while deadpan and grainyhead contained only one site (Fig. 2D). These analyses suggest that neuroblast-enriched miRNAs compose a regulatory module (Fig. 2E) that redundantly targets and downregulates most, if not all, key genes involved in initiating neuronal differentiation after asymmetric division, as well as master regulators of temporal identity.

Currently, few of these newly identified interactions have been experimentally validated (green lines in Fig. 2E). To exemplify their functionality, we focused on the miRNA showing strongest differential expression between neuroblasts and neurons, miR-1-3p (Fig.2B). While this miRNA has important regulatory roles in *Drosophila* and mammalian heart and muscles^42–45^, its expression has never been reported in the larval CNS. MiR-1-3p is predicted to target prospero, the best studied neurogenic factor in the fly, as well as three other mRNAs of the neurogenic module (nerfin-1, Eip93F, mamo; Fig. 2E). In the *Drosophila* late larval CNS, *prospero* is highly expressed in the Ganglion Mother Cell (GMC) and maturing neurons where it promotes differentiation and prevents reversion into neuroblast-like cells^46^ (Fig. 3A).

**Figure 3:**
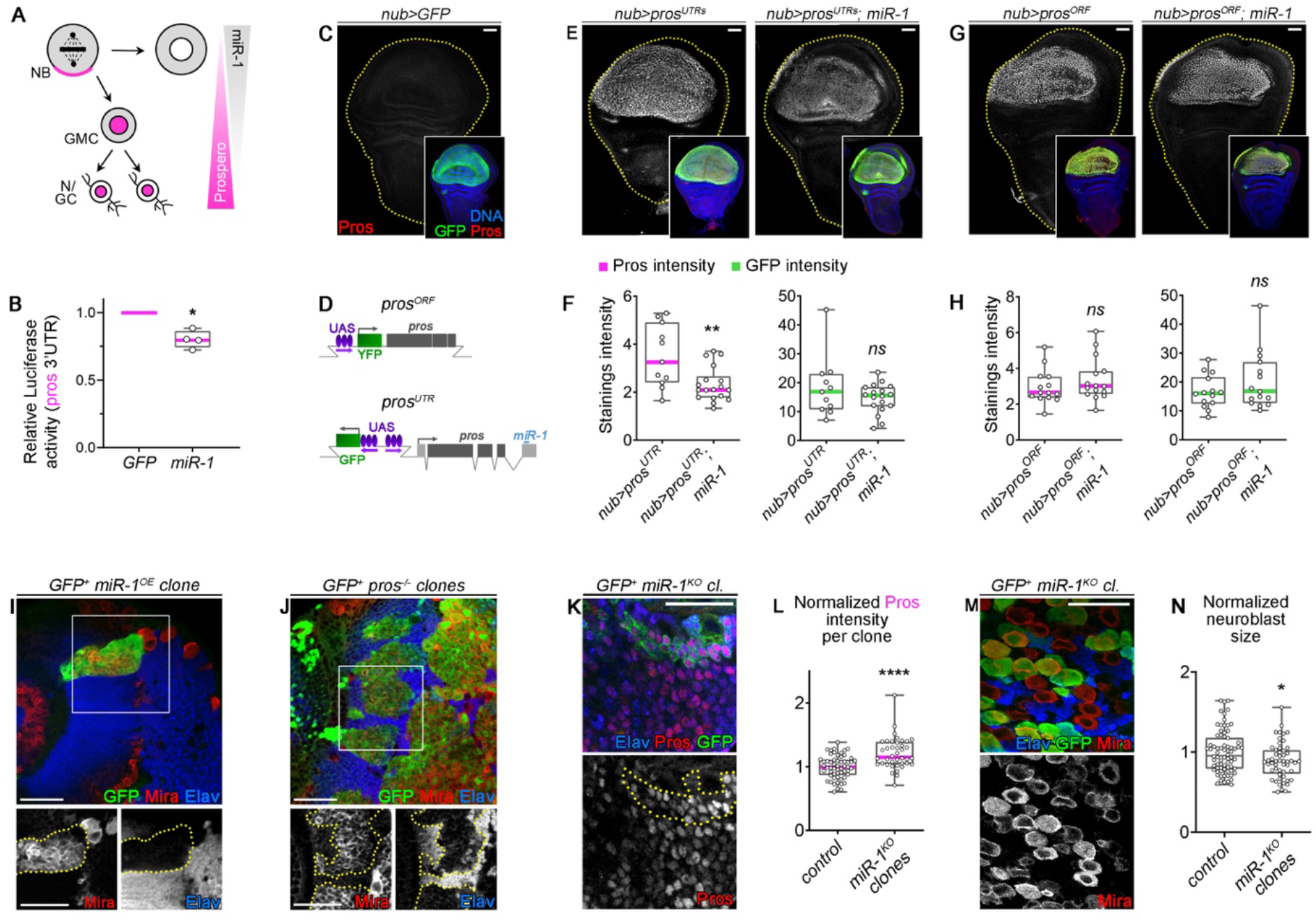
prospero mRNA is directly targeted by miR-1. **A)** Schematic representation of miR-1 and *prospero* expression during neurogenesis (NB: neuroblast, GMC: ganglion mother cell, N/GC: neuron/glial cell). **B)** Luciferase assay performed on HEK293 cells co-transfected with a luciferase reporter vector harboring the prospero 3’UTR together with either a GFP plasmid or a pri-miR-1 expressing plasmid. This assay showed a reduced luciferase activity upon miR-1 expression (n=4 independent experiments. p=0.02). **C)** Expression of GFP in the pouch of a *nub-GAL4, UAS-GFP* (*nub>GFP*) wing imaginal disc. Imaginal discs do not express endogenous *prospero* as shown by immunostaining against Prospero (Pros) (red). **D)** Schematic representation of the *UAS-YFP-prospero-ORF* (*pros^ORF^*) and *UAS-prospero-UTR*s (*pros^UTRs^*) transgenes. **E)** *nub-GAL4* drives *pros^ORF^* expression, alone or together with *miR-1,* in the wing pouch (labeled by YFP). **F)** Relative anti-Pros and anti-GFP immunostaining intensities in the wing pouch in *nub>pros^ORF^* (n= 14 wing discs, m= 2.96 ± 0.25 and m= 16.63 ± 1.52, respectively) and *nub>pros^ORF^*; miR-1-3p (n= 15 wing discs, m= 3.36 ± 0.31 and m= 20.27 ± 2.53, respectively). *p* = 0.234 and *p* = 0.451, respectively. Intensity of immunostaining for YFP and Pros are the same with or without *miR-1* co-expression. **G)** *nub-GAL4* drives *GFP* and *pros^UTRs^* alone or together with *miR-1* in the wing pouch. **H)** Relative anti-Prospero and anti-GFP staining intensities in the wing pouch in *nub>pros^UTRs^*(n= 11 wing discs, m= 3.50 ± 0.39 and m= 18.27 ± 3.15, respectively) and *nub>pros^UTRs^*; miR-1 (n= 18 wing discs, m= 2.32 ± 0.17 and m= 14.58 ± 1.20, respectively). *p* = 0.008 and *p* = 0.550, respectively. Intensity of immunostaining is reduced for Pros but not for GFP upon *miR-1-3p* co-expression indicating that miR-1-3p inhibits prospero *via* the 3’UTR. **I)** GFP+ neuroblast clones misexpressing *miR-1* in the OL. Neuroblasts and neurons are labeled by anti-Mira and anti-Elav immunostaining respectively. GFP+ clones are delineated by yellow dots. **J)** GFP+ *pros^-/-^* mutant clones in the OL immunostained with GFP in green, Mira in red and Elav in blue. Clones are delineated in yellow. Both *miR-1* overexpression and *pros* loss-of-function lead to neuroblast amplification at the expense of neurons. **K)** GFP+ *miR-1-3p^KO^* clones, stained with GFP in green, Elav in blue and Prospero in red. Clones are delineated in yellow. **L)** Normalized anti-Pros immunostaining intensity in control *wt* clones (n = 57 clones, 2 CNS, m = 0.99 ± 0.02) and in *miR-1-3p^KO^*clones (n = 40 clones, 3 CNS, m = 1.21 ± 0.04). *p* = 9.42 x 10^-6^. miR-1 knock-out leads to an increase in *pros* expression. **M)** GFP+ *miR-1-3p^KO^*clones stained with GFP in green, Mira in red and Elav in blue. Knock-out of *miR-1-3p* does not lead to precocious neuroblast differentiation. **N)** Normalized neuroblast area in control and in *miR-1-3p^KO^* clones. Neuroblast area is in average smaller in the *miR-1-3p^KO^* (n = 52 clones, 2 VNC+CB, m = 0.89 ± 0.03) condition than in the control condition (n = 66 clones, 5 VNC+CB, m = 1 ± 0.02) (*p* = 0.031). The scale bars represent 30 µm.

We investigated the molecular interactions of miR-1-3p with the prospero mRNA. *In vitro*, luciferase assays revealed that presence of the prospero 3’UTR significantly reduced expression upon co-expression of miR-1 (Fig.3B). We then used the wing imaginal disc of late larvae to test for the effect of miR-1 activity on prospero *in vivo*. *prospero* is not normally expressed in wing discs (Fig. 3C). We took advantage of a transgenic fly line with a *UAS* sequence inserted upstream of the *prospero* gene (Fig. 3D) to trigger mis-expression of endogenous *prospero* including its 3’ untranslated region (UTR) in the wing imaginal disc using the *nubbin-GAL4* driver line. This led to ectopic Prospero expression which was reduced when miR-1 was co-expressed (Fig. 3E,F). In contrast, mis-expression of a *UAS-prospero* transgene containing only the open reading frame (ORF), but lacking the 3’UTR, was insensitive to miR-1 (Fig. 3G,H). These results strongly suggest that the prospero mRNA is a direct target of miR-1-3p *in vitro* and *in vivo*. Moreover, over-expression of miR-1 in neuroblasts of the OL led to massive amplification of Mira positive neuroblasts at the expense of neuronal differentiation (Fig.3I), reminiscent of the well-known *prospero* loss-of-function phenotype^17,18,47^ and consistent with *prospero* silencing (Fig.3J). Comparably, overexpression of miR-9c, another neuroblast-enriched miRNA (Fig.2B) known to target another gene of the neurogenic module, *nerfin-1*^48^, led to clones containing a mix of Mira-positive neuroblasts and Elav-positive neurons, consistent with nerfin-1 inhibition^19,49^ (Fig.S3A,B).

Then we investigated the consequences of miR-1-3p and miR-9-5p loss-of-function. MiR-1 mutant clones in the VNC showed small but significant increases of Prospero in the neuronal progeny (Fig.3K,L), while clonal expression of a miR-9-5p sponge led to a minor but detectable increase in Nerfin-1 (Fig.S3C,D), in agreement with the predicted functional interactions.

Wild type neuroblasts differentiate during metamorphosis after a terminal symmetric division, following a period of progressive shrinking^28,50^. Ectopic expression of *prospero* or *nerfin-1* in larval neuroblasts causes precocious differentiation in larvae^19,28^. However, neuroblasts in clones were maintained in the VNC in miR-1 mutants (Fig.3M) or after expression of the miR-9-5p sponge (Fig.S3E), indicating that deregulation of their targets was insufficient to induce full differentiation. In the absence of strong defects, we investigated the presence of subtler signs of stemness exhaustion such as reduction in cell size^28,39,50,51^. Measuring neuroblast diameter revealed a small but significant (P= 0.003) difference in this parameter in miR-1 mutant flies (Fig. 3N). Thus, miR-1 and miR-9 over-expression efficiently downregulates two master regulators of *Drosophila* neurogenesis in the progeny of neuroblasts and inhibits their proper differentiation. However, loss-of-function is insufficient to trigger neuroblast termination, although it can induce minor signs of differentiation in the case of miR-1.

### miRNA of the neurogenic module work in concert

Our bioinformatic analyses identified a large set of neuroblast-enriched miRNAs, all predicted to redundantly target key regulators of the neuroblast to neuron transition (Fig. 2E). This points to a scenario in which, at physiological expression levels, the cooperative action of multiple miRNAs is required to achieve functionally relevant levels of regulation. We aimed at testing this possibility by comparing the effect of knocking down either single or multiple miRNAs on neuroblasts. To structure the approach, we sought to identify functional sub-groups among the differentially regulated miRNAs. Hierarchical clustering based on their predicted target genes revealed the presence of three main groups (Fig. 4A). Three miRNAs (miR-306-5p, miR-988-3p and miR-995-3p), which are predicted to target only few genes controlling neuroblast to neuron transition (Fig 2C), clearly clustered separately and were not considered in further analyses. The remaining 10 miRNAs formed two distinct clusters based on their target preferences, with miR-1-3p and miR-9-5p as members of cluster 1 (Fig. 4A). miR-11-5p and miR-9c-3p were expressed at low levels, and therefore were not considered for further experiments. Next, we compared the effect of knockdown of either individual or entire clusters of the identified miRNAs on neurogenesis. To study individual miRNAs we used either established transgenic sponge lines^52^ or, for miR-1-3p, miR-34-3p as well as the related miR-279-3p and miR-996-3p, we generated new fly lines (Fig. S4A; Table S1).

**Figure 4:**
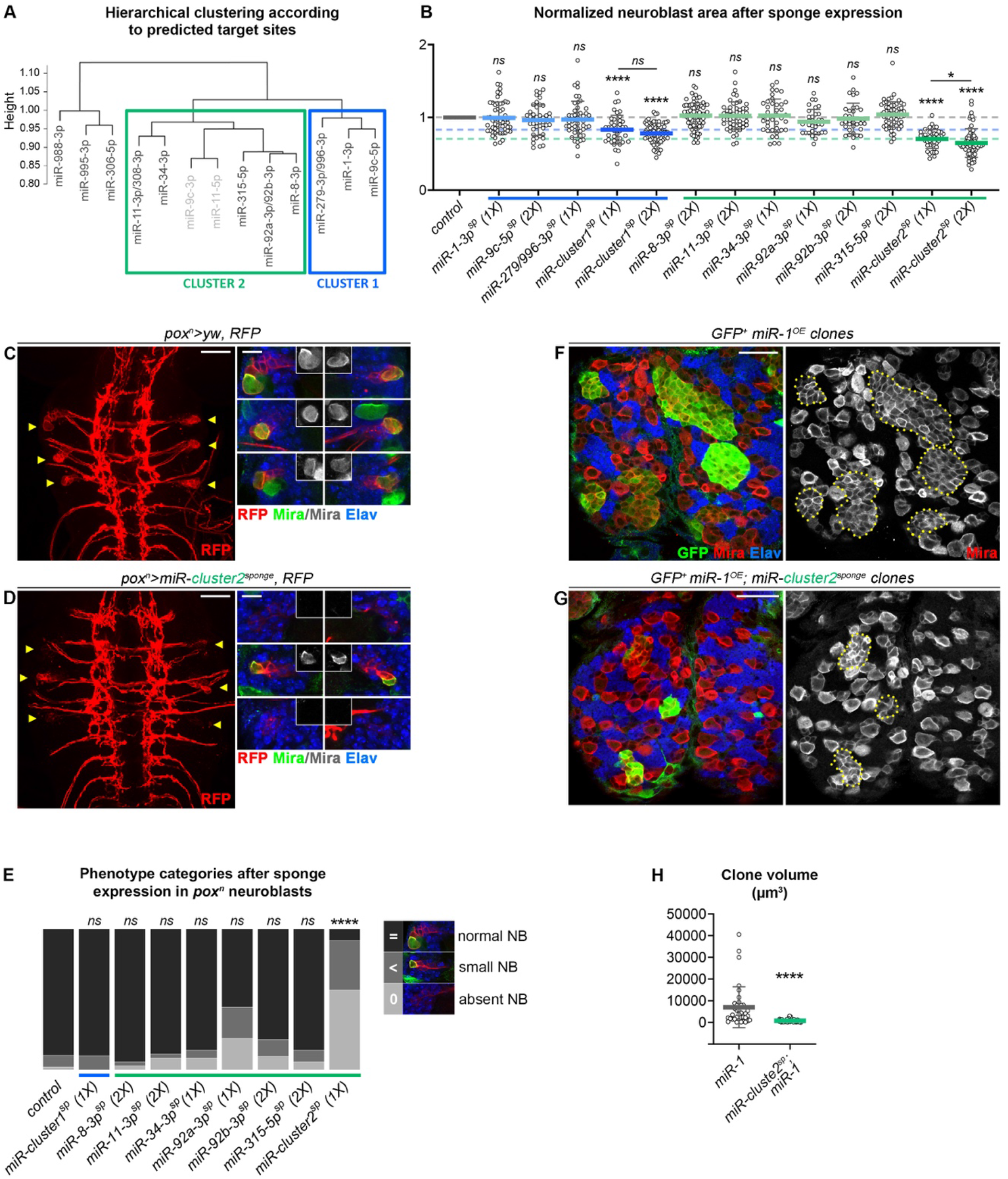
miRNAs cooperate to regulate neuronal differentiation. **A)** Hierarchical clustering of neuroblast-enriched miRNAs based on the similarity of their predicted mRNA targets. MiRNAs in light gray (miR-11-5p and miR-9c-3p), expressed at much lower level, were not targeted by the multi-sponge constructs. **B)** Comparison of normalized neuroblast area in VNC clones expressing sponges for individual miRNAs or cluster-1 and cluster-2: control (n = 66 clones, 5 CNS, m = 1 ± 0.02), *miR-1-3p^sponge^* (1X) (n = 58 clones, 3 CNS, m = 0.99 ± 0.03), *miR-9c-5p^sponge^* (2X) (n = 43 clones, 2 CNS, m = 0.96 ± 0.03), *miR-279/996-3p^sponge^* (1X) (n = 48 clones, 3 CNS, m = 0.97 ± 0.04), *miR-cluster1^sponge^*(1X) (n = 41 clones, 3 CNS, m = 0.83 ± 0.03), *miR-cluster1^sponge^*(2X) (n = 70 clones, 3 CNS, m = 0.78 ± 0.02), *miR-8-3p^sponge^* (2X) (n = 73 clones, 2 CNS, m = 1.02 ± 0.02), *miR-11-3p^sponge^* (2X) (n = 54 clones, 3 CNS, m = 1.02 ± 0.02), *miR-34-3p^sponge^* (1X) (n = 40 clones, 3 CNS, m = 1.02 ± 0.03), *miR-92a-3p^sponge^* (1X) (n = 33 clones, 3 CNS, m = 0.94 ± 0.03), *miR-92b-3p^sponge^* (2X) (n = 35 clones, 4 CNS, m = 0.98 ± 0.02), *miR-315-5p^sponge^* (2X) (n = 60 clones, 4 CNS, m = 1.04 ± 0.02), *miR-cluster2^sponge^* (1X) (n = 41 clones, 4 CNS, m = 0.70 ± 0.02), *miR-cluster2^sponge^* (2X) (n = 69 clones, 5 CNS, m = 0.65 ± 0.02). The p-values issued from comparison of each sponge construct with control are: *p* = 0.429, *p* = 0.256, *p* = 0.226, *p* = 2.59 x 10^-5^, *p* = 3.89 x 10^-11^, *p* = 0.300, *p* = 0.727, *p* = 0.728, *p* = 0.264, *p* = 0.370, *p* = 0.109, *p* = 4.90 x 10^-12^ and *p* = 1.04 x 10^-20^, respectively. Thus, only miR-cluster 1 and 2 sponge constructs showed significant differences with control. Moreover, *miR-cluster2^sponge^*(1X) clones showed a significantly larger neuroblast size than *miR-cluster2^sponge^* (2X) (*p* = 0.011). 1X or 2X indicates the number of copies of the sponge transgene carried by the fly. **C)** Control VNC expressing RFP under the control of *pox^n^-GAL4*. Yellow arrowheads highlight pox^n^ lineages in late larvae, which are shown enlarged on the right. Each lineage contains an RFP+ neuroblast (Mira+) and several RFP+ progeny (GMCs and neurons (Elav+)). **D)** Most *pox^n^* lineages (yellow arrowheads) expressing the *miR-cluster2^sponge^* transgene (*pox^n^-GAL4; UAS-miR-cluster2^sponge^*) lose neuroblasts **E)** Distribution of neuroblast phenotypes after sponge expression using *pox^n^-GAL4* : control (n = 49 NBs), *miR-cluster1^sponge^* (1X) (n = 30 NBs), *miR-8-3p^sponge^*(2X) (n = 36 NBs), *miR-11-3p^sponge^* (2X) (n = 36 NBs), *miR-34-3p^sponge^* (1X) (n = 36NBs), *miR-92a-3p^sponge^* (1X) (n = 42 NBs), *miR-92b-3p^sponge^* (2X) (n = 42 NBs), *miR-315-5p^sponge^* (2X) (n = 36 NBs), and *miR-cluster2^sponge^* (1X) (n = 60 NBs). The p-adjusted-values obtained after pairwise comparison of each sponge construct as indicated from letf to right with control (no sponge) were: *p* = 1, *p* = 1, *p* = 1, *p* = 1, *p* = 0.0673, p=1, *p* = 1 and *p* = 5.31 x 10^-19^ respectively. *p* = 0,00000418 compares the clones in *miR-92a-3p^sponge^* (1X) sponge condition with the *miR-cluster2^sponge^* (1X) condition. **F,G)** GFP-labeled clones misexpressing *miR-1* only (F) or *miR-1 plus cluster-2^sponge^* (G) stained with GFP in green, Mira in red and Elav in blue. Clones are delineated in yellow. **H)** Mean clone volumes quantified in clones over-expressing *miR-1* (n = 36 clones, 3 CNS, m = 6989 µm^3^ ± 870 µm^3^) or *miR-1; miR-cluster-2^sponge^* (n = 45 clones, 3 CNS, m = 796 µm^3^ ± 58 µm^3^). *p* = 1.11 x 10^-8^. The scale bars represent 30 µm, except for the blow-ups in C and D where the scale bars represent 10 µm.

Two multi-sponge fly lines based on assembled sponge constructs targeting all miRNA of either cluster 1 (miR-1-3p, miR-9c-5p and miR-279-3p/996-3p) or cluster 2 (miR-8-3p, miR-11-5p, miR-92a/b-3p, miR-315-5p, miR-34-3p) were also generated (Fig S4A). In contrast to genetic miR-1^KO^ (Fig. 3M), we found that sponge mediated knock-down of this miRNA, or of other individual miRNAs in the module, had never a statistically significant effect on neuroblast size (Fig. 4B), in agreement with the expected incomplete inhibition induced by sponges. However, expression of either multi-sponge 1 or multi-sponge 2 significantly reduced neuroblast size in a dose dependent manner (Fig. 4B, S4B). Thus, the effect of cluster knockdown on neuroblast size exceeded that of any sponge targeting an individual miRNA. These phenotypes are consistent with the de-repression of the neural differentiation gene network composed by the predicted miRNA targets^51^. We conclude that miRNAs act additively, or even cooperatively, in this context.

Next, we investigated the impact of the neurogenic miRNA module on a specific subset of neuroblasts and their offspring. The *pox^n^* driver has been previously used to study the neurogenic properties of six defined neuroblasts in the VNC^47,53,54^ (Fig. 4C). Expression of the multi-sponge construct targeting the entire cluster 1 or the individual miRNAs of cluster 2 did not induce statistically significant phenotypes (Fig. 4D,E). However, multi-sponge 2 expression promoted significant shrinkage and often caused precocious elimination of *pox^n^*-positive neuroblasts in late larvae (Fig. 4D,E). This strong phenotype highlights that some neuroblasts are particularly sensitive to the knockdown of the miRNA module and suggests that they require this module to maintain stemness potential throughout development.

To further investigate the cooperative aspect of this regulatory system we combined gain- and loss-of-function approaches. As shown above, miR-1 over-expression efficiently silenced prospero mRNA in neuroblasts and GMCs and induced their amplification (Fig. 4F). Interestingly, co-overexpression of miR-1 with the multi-sponge against cluster 2 miRNAs, led to strong reduction of neuroblast numbers per clone (Fig. 4F-H), showing that neuroblast amplification was partially suppressed. Thus, increased function of one miRNA of the module can be counterbalanced by lowering the efficiency of other members.

## Discussion

A given miRNA is predicted to bind hundreds of mRNAs and gain-of-function studies generally validate these predicted interactions, showing repression of targets and mainly strong phenotypic changes ^55,56^. In contrast, loss of function experiments lead in general to a very limited increase in target gene expression^4,5^ and induce only mild, often non-detectable, phenotypes^6,7,57^. Based on these observations, it is considered that under physiological conditions miRNAs act as rheostats of biological processes, ensuring the robustness of a specific molecular pathway^58,59^. However, such a limited role as simple balancer of protein production appears contradictory to the finding that impairment of the entire miRNA pathway, for example through constitutive or conditional inactivation of dicer results in general in the rapid induction of strong phenotypes in the affected tissue, pointing to a much more active regulatory role^60,61^. In light of these seemingly opposing features, additive or synergistic actions of groups of miRNAs to control a biological process represent an attractive solution. Indeed, in this scenario a significant reduction of target gene expression is only achieved through the joined action of several miRNAs on a specific target mRNA, while a given biological process is regulated through the joint quantitative regulation of expression of entire groups of genes. Here, we tested this hypothesis in the context of brain development in Drosophila melanogaster, where neuroblasts undergo asymmetric divisions, allowing production of neurons and glia as they cycle throughout development. This stereotypic process is highly controlled to ensure that neurons and glia are produced with the correct identity and in correct numbers and miRNAs represent attractive candidates to play key roles in these developmental processes.

As an entry point, we developed in vivo AGO-APP to isolate microRNAs from neuroblasts, neurons and glia. Using transgenic flies, we demonstrate that this approach allows the isolation and analysis of AGO-bound miRNAs with highest precision. Indeed, in all analyzed cases the expression levels and dynamics observed in the AGO-based sequencing approach were confirmed by independent findings in *Drosophila* reporter lines that allow to visualize the expression of specific miRNAs. Moreover, AGO-miRNAs that are only present in small subpopulations, like for example miR-10, are reliably detected in a strong background of negative cells, showing that APP is highly sensitive. Besides its precision and sensitivity, AGO-APP has other experimental advantages. First, AGO-APP is a benchtop approach through which cell type specific miRNAs can be isolated without the use of complex cell isolation approaches like, for example Fluorescent Activated Cell Sorting. Second, AGO-APP binds all AGO proteins with high affinity, allowing the isolation of the entire spectrum of miRNAs from all species. Third, non-AGO bound miRNAs and other small RNAs are excluded by the approach, lowering complexity and allowing to focus only on the RNA population of interest, such that are actively involved in inhibition.

Moreover, knowing that the T6B peptide can be attached to proteins showing particular subcellular localizations, like the nucleus, cytoplasm or synapses, it may be possible in the future to study the expression of AGO-bound RNAs in different cellular compartments.

Using a non-biased statistical approach, we identified a group of neuroblast-enriched miRNAs that are predicted to target almost all factors known to control early steps of the neuroblast-to-neuron transition, pointing to the existence of a regulatory module that cooperatively maintains NSC status versus differentiation. Overexpression of individual miRNAs of the group, namely miR-1 and miR-9, confirmed their capacity to induce dominant differentiation defects in neuroblast progeny, phenotypes mimicking the silencing of the key differentiation genes that are their targets. Thus, it is important that the expression level of each miRNA is tightly regulated along neuroblast lineages. Interestingly, functional analyses based on sponge mediated knockdown demonstrated that loss of individual miRNAs has no or little detectable influence on neuroblast status suggesting that, in normal conditions, miRNAs tend to be dispensable when depleted individually. However, simultaneous downregulation of multiple miRNAs of the module induces a significant biological response in the form of precocious stemness exhaustion. Altogether these results are fully consistent with a situation where a set of specific miRNAs, expressed at low levels, is actively protecting neuroblasts against precocious differentiation through a sum of weak and redundant inhibitory effects on neurogenic factors. Interestingly, this mechanism also provides a strategy to refine the number of silenced mRNA targets to the ones containing several binding sites for the miRNAs that constitute the module. In this sense, it is striking that all iconic neural differentiation factors, and many temporal factors, are among the top 1.5% of mRNAs most repeatedly targeted by the neuroblast-specific miRNA module. This also shows that beyond defining a set of functionally significant target genes, miRNA modules may be used to target genes involved in a common biological process. Thus, our study provides strong evidence that the cooperative action of miRNAs on sets of target genes confers both, the specificity of the genes to be silenced and the efficiency of silencing.

Finally, such cooperation between miRNAs of the same module is also supported by the finding that the dominant NB overproliferation phenotype caused by over-expression of a single miRNA, miR-1, depends on the expression of the other miRNAs of the module. Indeed, miR-1, overproliferation can be efficiently inhibited through the partial knockdown of other members of the module. Given that miRNA overexpression has been implicated in numerous cancers^62^, and in light of the possibility of using sponges to partially inhibit multiple miRNAs, identifying miRNA modules acting in concerts with oncogenic miRNA could represent an opportunity for developing new therapeutic approaches.

## Acknowledgements

The Cremer lab has been supported by the ANR (MicroRNAct, ANR-17-CE16-0025; Uncoding, ANR-21-CE16-0034; Miniature, NR-21-CE13-0003; Goligo, ANR-21-CE16-0023) and the Fondation pour la Recherche Médicale (Program Equipe FRM). Andrea Erni received a postdoc fellowship from the Swiss National Funds; Surbhi received a PhD fellowship from Neuroschool Marseille. The Maurange lab was funded by ANR (Miniature, NR-21-CE13-0003) and the Ligue Contre le Cancer (programme Equipe Labellisée).

We also thank the local PiCSL-FBI core facility [Institut de Biologie du Développement de Marseille (IBDM), Aix-Marseille University] supported by the French National Research Agency through the “Investments for the Future” program (France-BioImaging, ANR-10-INBS-04) as well as the IBDM animal facilities.

## Figure Legends

**Figure S1:**
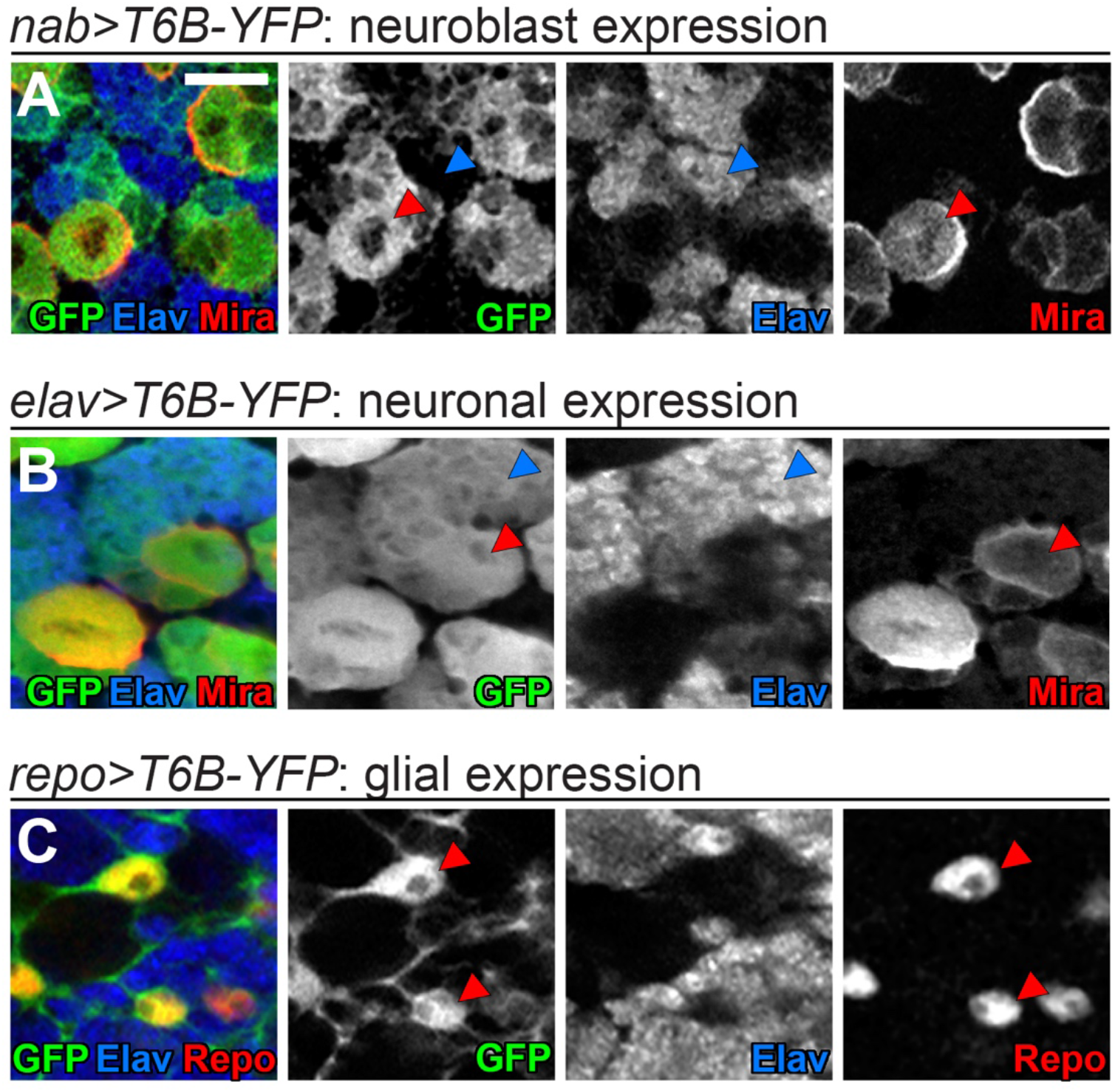
Immunostaining validation of T6B expression pattern in the CNS of the different *Drosophila* lines used to perform cell-type specific Ago-APP. **A)** *UAS-T6B-FYH* transgene expression, revealed by an anti-GFP immunostaining (green), driven by *nab-GAL4*, strongly labels Mira-positive neuroblasts in red (red arrowhead) as well as GMCs and a few recently born Elav-positive neurons in blue due to protein perdurance after asymmetric neuroblast division. However, GFP is absent from mature elav-positive neurons (blue arrowhead). **B)** *UAS-T6B-FYH* transgene expression, revealed by an anti-GFP immunostaining (green). Staining shows that *elav-GAL4* is active in neuroblasts labeled with an anti-Miranda (Mira) antibody (red arrowheads) and all neurons labeled with an anti-Elav antibody (blue arrowhead) **C)** *UAS-T6B-FYH* transgene expression, revealed by an anti-GFP immunostaining (green), using the glia specific driver *repo-GAL4*. GFP is expressed in Repo-positive glia (red) (red arrowheads) but absent in Elav-positive neurons (blue). Scale bars represent 10 µm.

**Figure S2:**
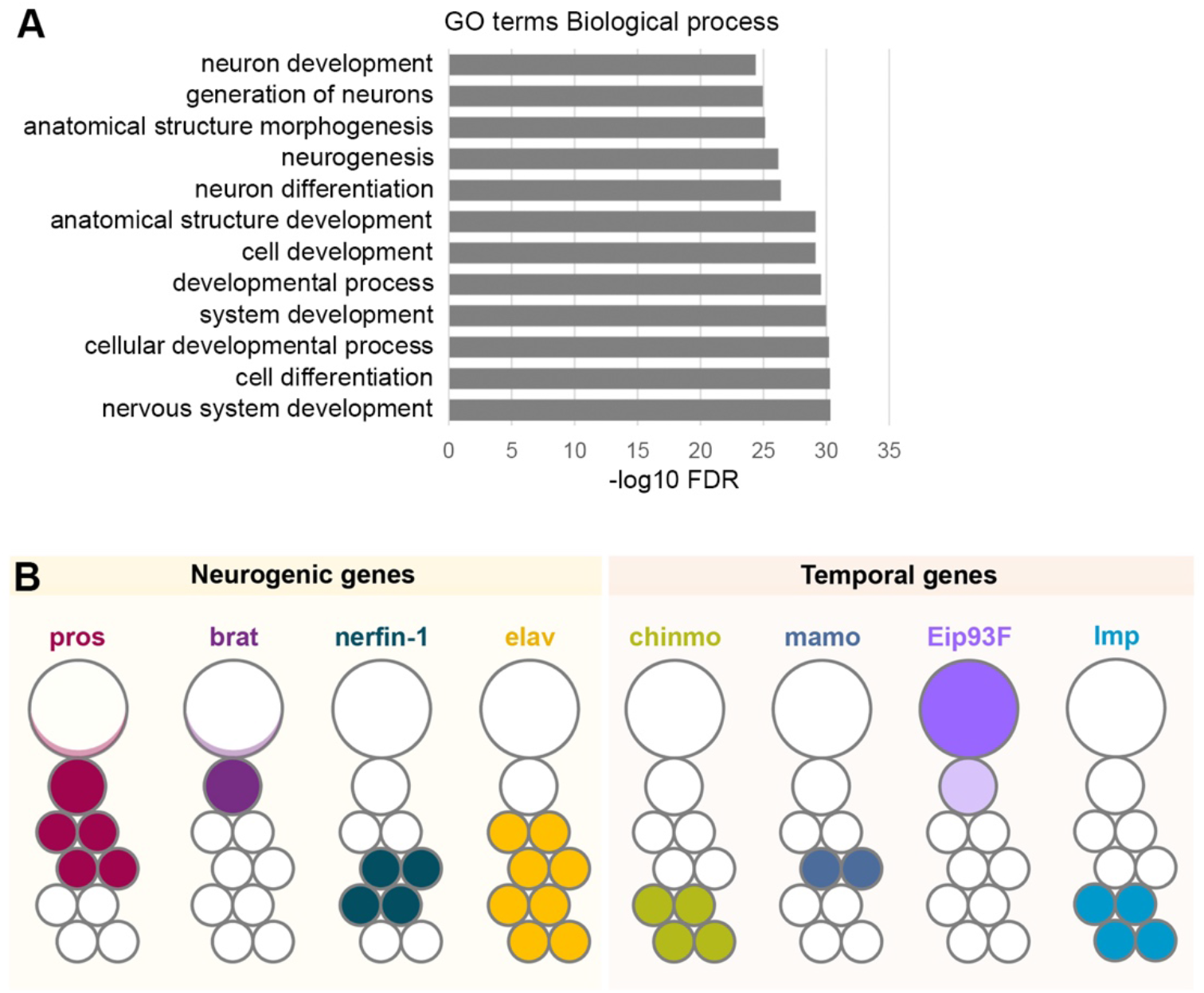
Neuroblast-enriched miRNAs target neuronal differentiation. **A)** List of the most significant terms issued from a GO analysis performed on the 227 predicted mRNAs coded by the drosophila genome predicted to be targeted at least 5 times by the module of neuroblast-specific miRNAs. Most of these terms are related with neurogenesis and neuron differentiation **B)** Schematic representation of expression pattern of iconic neurogenic and temporal genes along a neuroblast lineage in late larvae. These genes are predicted to exhibit multiple binding sites for the neuroblast-enriched miRNA module, as depicted in Figure 2E.

**Figure S3:**
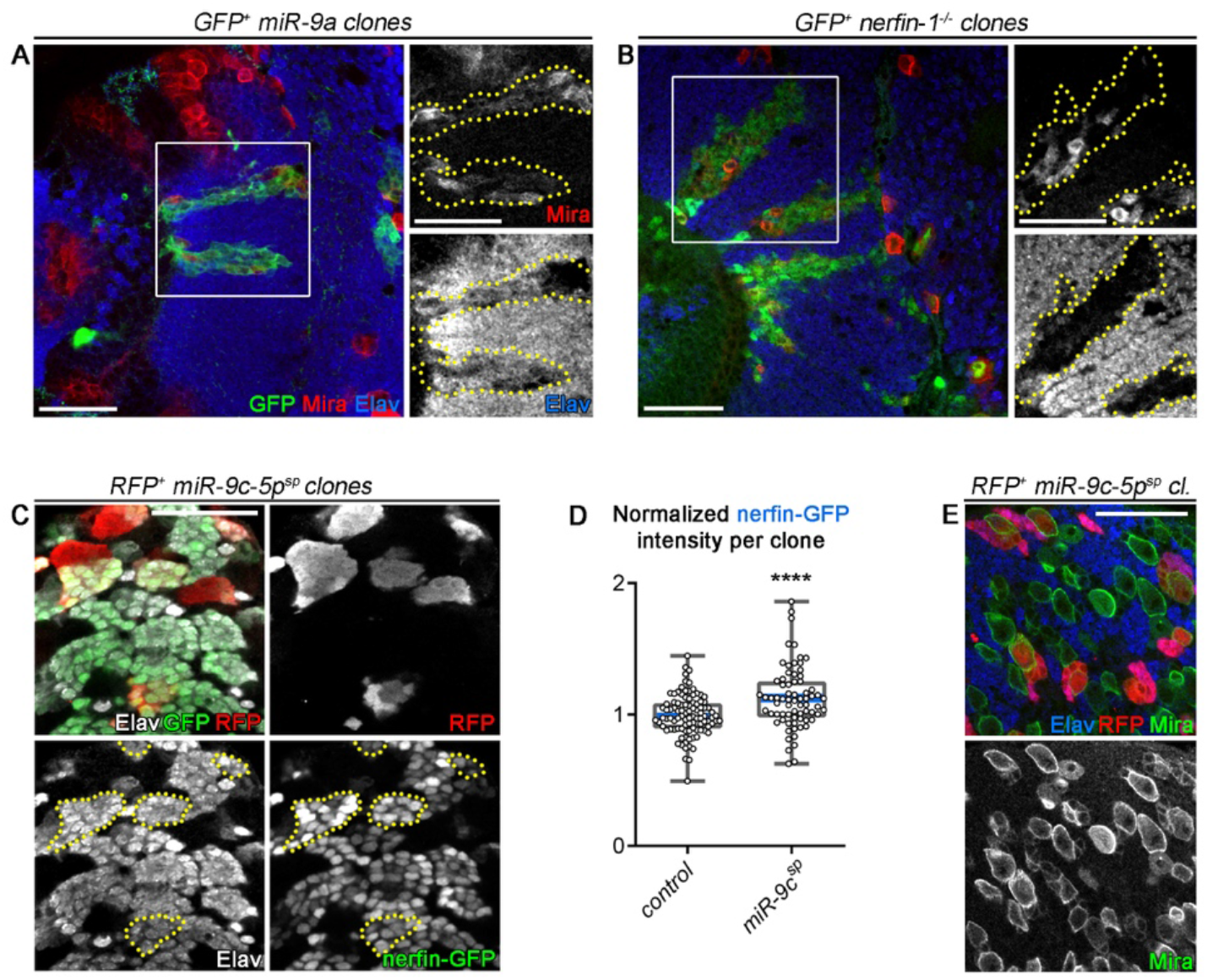
nerfin-1 is inhibited by miR-9. **A,B)** GFP+ neuroblast clones over-expressing *miR-9a* (A) and GFP+ neuroblast *nerfin-1^-/-^* clones (B) in the OL immunostained against GFP in green, Mira in red and Elav in blue. Clones are delineated in yellow. Both *miR-9a* over-expression and *nerfin-1* loss of function leads to neuroblast amplification. Note that clones are composed of a mix of neuroblasts and neurons, unlike in *miR-1^OE^* and *pros^-/-^* clones (Fig. 3I). **C)** RFP-labeled clones expressing *miR-9c-5p^sponge^*stained with RFP in red, Elav in white and nerfin-1-GFP in green. Clones are delineated in yellow. **D)** Normalized nerfin-GFP intensity in control *wt* clones (n = 98 clones, 4 CNS, m = 0.99 ± 0.02) and in clones expressing *miR-9c-5p^sponge^*(2X) (n = 71 clones, 5 CNS, m = 1.13 ± 0.03). *p* = 1.08 x 10^-4^. **E)** RFP+ clones expressing *miR-9c-5p^sponge^* stained with RFP in red, Mira in green and Elav in blue. Knock-down of *miR-9c-5p* does not lead to neuroblast loss.

**Figure S4:**
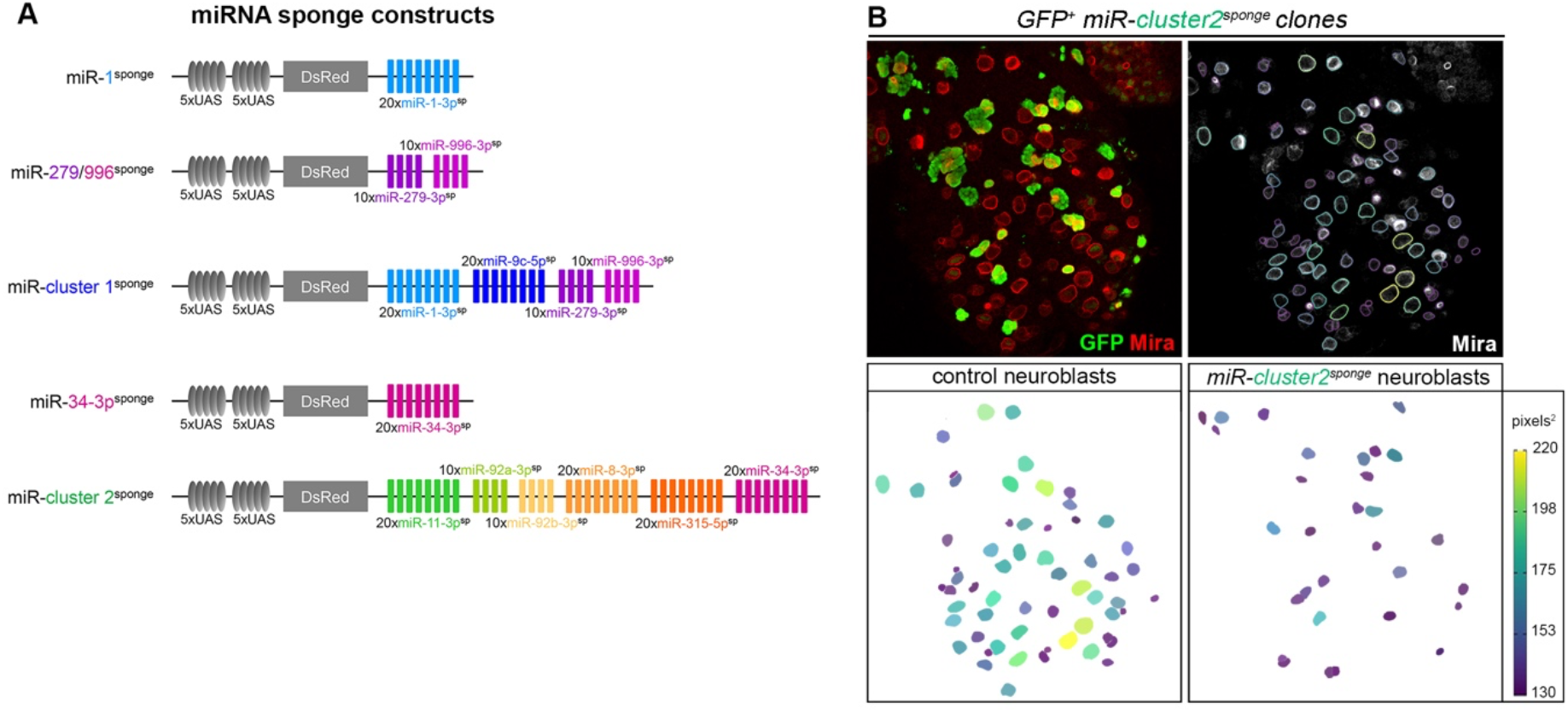
Sponge-mediated knockdown of miRNAs. **A)** Sponge constructs made in this study. They consist of *20* miRNA binding sites with mismatches at positions 9–12 placed in the 3ʹ-untranslated region of DsRed under the control of 10 tunable UAS binding sites. Transgene integration has been done in one copy at the landing site attp40 on the second chromosome for *miR-1-3p^sponge^*, *miR-279/996-3p^sponge^* and *miR-cluster1^sponge^* and at the landing site attp2 on the third chromosome for *miR-34-3p^sponge^* and *miR-cluster2^sponge^*. **B)** GFP-labeled clones in the late larval CNS misexpressing *miR-cluster2^sponge^*. All neuroblasts are color-coded relative to size (in pixels^2^). Control neuroblasts (non-GFP labelled) and neuroblasts expressing the *miR-cluster2^sponge^* (GFP-positive) are shown separately, showing that NBs misexpressing *miR-cluster2^sponge^* are smaller.

**Table S1:**
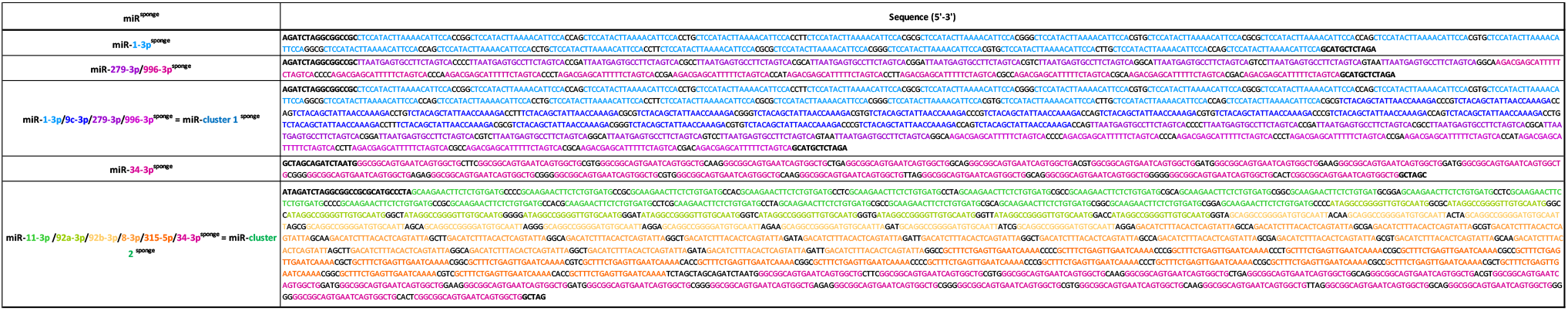
Sequences used for sponge constructs generated in this study.

## Material and Methods

### Fly lines

Drosophila stocks were maintained at room temperature on standard medium (8% cornmeal, 8% yeast, 1% agar). Experiments were performed at 29 °C. Crosses to yw (Bloomington #1495) line are used as controls. For generating *pros^-/-^*, *nerfin-1^-/-^* and *miR-1^KO^* clones, we used the Mosaic analysis with a repressible cell marker (MARCM) technique^63^. The following MARCM stocks were used : tub-GAL4, UAS-nGFP-myc, hs-FLP; FRT82B, tub-GAL80/TM6c, FRT82B, pros^17^/TM6b (from Bloomington #5458), tub-GAL4, UAS-nGFP-my,c hsFLP122; tub-GAL80^LL9^, FRT2A/TM6b and Df(3L)nerfin-1^159^, FRT2A/TM6. For generating clones mis-expressing miRs or sponges, we used the Flp-out (Flp^out^) technique. Flp^out^ clones were generated using hs-FLP; Act5c>CD2>GAL4, UAS-GFP or hs-FLP; Act5c>CD2>GAL4, UAS-RFP/TM3 (from Bloomington #7 and #30558). The GAL4 lines used were the following: nab-GAL4 (Kyoto DGRC #6190), repo-GAL4 (Bloomington #7415), elav-GAL4 (Bloomington #8765), nubbin-GAL4 (Bloomington #86108), miR-10^KO^-GAL4 (Bloomington #58880). The UAS lines used were: UAS-LUC-miR-1/TM3 (Bloomington #41125), UAS-LUC-miR-8 (Bloomington #41176), UAS-LUC-miR-9a (Bloomington #41138), same result than with UAS-LUC-miR-9c (Bloomington #41139), UAS-DsRed2-miR-11 (Bloomington #59864), UAS-LUC-miR-92a (Bloomington #41153), UAS-LUC-miR-92b (Bloomington #41175), UAS-LUC-miR-279 (Bloomington #41147), UAS-miR-315 (FlyORF #F002078), UAS-YFP-prospero (prospero cDNA, called here pros^ORF^, from Andrea Brand); UAS-prospero (transgene inserted in 5’ of prospero, called here pros^UTRs^, Kyoto DGRC #201789), UAS-mCherry-miR-8^sponge^ (Bloomington #61374), UAS-mCherry-miR-9c^sponge^ (Bloomington #61376), UAS-mCherry-miR-11^sponge^ (Bloomington #61378), UAS-mCherry-miR-92a^sponge^ (Bloomington #41153), UAS-mCherry-miR-92b^sponge^ (Bloomington #41175), UAS-mCherry-miR-315^sponge^ (Bloomington #61432). For each of these miR^sponge^ lines, 2 copies -2X-of the miR^sponge^ transgene are present, except when indicated, on both the attp40 and attp2 landing sites. tub-GAL80^ts^ (Bloomington #7017 and #7019) was used to avoid premature GAL4 expression when at restrictive temperature (18°C). UAS-nlsGFP (Bloomington #4776) or UAS-mCD8ChRFP (Bloomington #27392) were used to follow the driver expression. The progeny of the crosses using the Flp-out technique was heat-shocked 1 hour at 37°C, raised at 29°C for 3 days and dissected at wandering L3 stage. The progeny of the crosses using the GAL4 technique was raised at 29°C (miRs expression experiments), or were maintained at 18°C, and switched to 29° one day before dissection. (miR-7)E>GFP^3^, 16.6 kb mir-279/996-GFP^4^, nerfin-1-GFP^5^ were used to monitor miR-7, miR-279/996 and nerfin-1 expression, respectively.

### Immunohistochemistry

Dissected tissues were fixed 5 to 15 minutes in 4 % formaldehyde/PBS depending on the primary antibody. Stainings were performed in 0.5 % triton/PBS with antibody incubations separated by several washes. Tissues were then transferred in Vectashield with or without DAPI for image acquisition. Primary antibodies used: chicken anti-GFP (1:1000, Aves #GFP-1020), rat anti-RFP (1:500, Chromotek #5F8), rabbit anti-RFP (1:500, Rockland #600-401-379), mouse anti-Miranda (1:20, A. Gould), rat anti-Dpn (1:50, abcam #195173), rat anti-Elav (1:50, DSHB #7E8A10), mouse anti-Elav (1:50, DSHB #9F8A9), mouse anti-Repo (1:200, DSHB #8D12) and mouse anti-Prospero (1:20, DSHB #MR1A). Adequate combinations of secondary antibodies (Jackson Immunodetect) were used to reveal expression patterns.

### Image processing

Confocal images were acquired on a Zeiss LSM 780 microscope (Zeiss, Oberkochen, Germany). FIJI and Photoshop were used to process confocal data. In each picture, the scale bar represents 30 µm.

To measure relative Pros and nerfin-GFP signal intensities in the CNS after immunostaing, for a given focal plane, the intensity of Pros (or GFP) staining in RFP-positive (i.e. clonal) neurons was divided by the mean intensity of Pros (or GFP) staining in surrounding RFP-negative (i.e. wild-type) neurons of the same focal plane. Both intensities were obtained with the “Mesure” plugin in FIJI.

To measure relative Pros and GFP intensities in the disc, for a given focal plane, the intensity of Pros (or GFP) staining in the cells of the wing pouch was divided by the intensity of Pros (or GFP) staining in the cells outside of the wing pouch delineated by morphological criteria revealed by Hoechst (not shown). Both intensities were obtained with the “Mesure” plugin in FIJI.

The relative area of each neuroblast within a clone expressing a sponge was the ratio between the area of the neuroblast (manually delimited using the Mira staining and measured using the “Mesure” plugin in FIJI) and the mean area of the wild-type surrounding NBs of the same focal plane.

### Statistical analysis

Statistical analyses were performed in R using rstatix package. We performed a Wilcoxon-Mann-Whitney test when comparing means of 2 independent groups. For comparing the repartition of categories between the independent multiple conditions of Fig 4E, we performed a Chi2 test followed by a p-value adjustment for multiple comparisons using the pairwise_chisq_gof_test function in rstatix. For Fig. 3B,F,H, L and Fig. S3E,H results are presented as boxes, extend from the 25^th^ to 75^th^ percentiles with the median, down to the minimum value, up to maximum value, and show all individual values as a point. For Fig.4B,H, results as scatter dot plots, and show all individual values as a point. For Fig.4E, each category is represented with bars in percentages. p-values are issued from comparison of each sponge construct with control. The sample size (n), the mean ± the standard error of the mean (m ± SEM), and the p-value are reported in the figure legends. ****p-value ≤ 0.0001, ***p-value ≤ 0.001, **p-value ≤ 0.01, *p-value ≤ 0.05 and ns p-value>0.05.

### Generation of *Drosophila* transgenic lines

Generation of the UAS-T6B lines: The T6B sequence fused to YFP,HA and FLAG was cloned downstream of the UAS sequence into the transformation vector pUASTattB^64^. The transgene was inserted into the attp2 landing site on chromosome III.

Generation of sponge lines: Sponge constructs were synthetized by GenScript Biotech Corporation (NL). Sponge constructs aimed at inhibiting a specific miRNA were generated using previously published guidelines^52^. Briefly, 20 repeats of a 21nt sequence fully complementary to the miRNA except mismatches at positions 9–12 of the miRNA, were assembled separated by variable four-nucleotide linkers, into a modified pWalium-ChtVis-Tomato vector^9^ (Addgene #67756). MiR-cluster1^sponge^ (miR-1-3p/9c-3p/279-3p/996-3p^sponge^) and miR-cluster2^sponge^ (miR-1-3p/9c-3p/279-3p/996-3p ^sponge^) were generated in the same modified pWalium-ChtVis-Tomato vector by simply assembling single sponge constructs. One copy (1X) of each sponge has been inserted into the attp40 (chromosome II) landing site for miR-1-3p^sponge^, miR-279-3p/996-3p^sponge^ and miR-cluster1^sponge^ or into the attp2 landing site (chromosome III) for miR-34-3p^sponge^ and miR-cluster2^sponge^.

The different genetic fly lines were generated by BestGene Inc. (http://www.thebestgene.com/).

### Generation of the anti-Miranda antibody

Anti-Miranda polyclonal antibody was generated by Genscript (https://www.genscript.com/). The epitope used for immunization is: mhhhhhhhsfskaklkrfndvdvaicgspaasnssagsagsatptassaaaapptvqperkeqiekffkdavrfassskeakefaipkedkk skglrlfrtpslpqrlrfrptpshtdtatgsgsgastaastplhsaattpvkeaksasrlkgkealqyeirhkneliesqlsqldvlrrhvdqlkeaeak lreehelatsktdrliealtsenlshkalneqmgqehadllerlaameqqlqqqhdeherqvealvaesealrlanellqtanedrqkveeqlq aqlsalqadvaqarehcsleqaktaenielvenlqktnaslladvvqlkqqieqdalsygqeakscqaeleclkverntlkndlankctlirslqd elldknceidahcdtirqlcreqarhteqqqavakvqqqvesdlesavereksywraeldkrqklaenelikielekqdvmvllettndmlrm rdeklqkceeqlrngidyyiqlsdalqqqlvqlkqdmaktitekynyqltltntratvnilmerlkksdadveqyraelesvqlakgaleqsylvlq adaeqlrqqltesqdalnalrsssqtlqseiansfqeridgdaqlahyhelrrkdetreaymvdmkkaldefatvlqfaqleldnkeqmlvkvr eeceqlklenialkskqpgsasllgtpgkanrsnttdlekiedllcdselrsdcekittwllnssdkcvrqdttseinellsagkssprpaprtpkap htprsprtphtprtprsaastpkktvlfagkenvpsppqkqvlkarni.

### Lysate preparation and Ago-APP

T6B expressing *Drosophila* CNS were isolated and fixed with 4% PFA for 10 minutes and subsequently quenched in a final concentration of 150 mM Glycine for 5 minutes, at room temperature each. After washing twice with ice-cold PBS, CNS were shock-frozen and stored at -70 °C until use. Ago-APP was performed according to previously published protocol^25^ with minor modifications. Tissue was lysed in 1ml lysis buffer (150 mM KCl, 25 mM Tris, pH 7.5, 2 mM EDTA, 0.5% Nonidet P-40; supplemented with 1 mM NaF, 1 mM DTT and 1 mM AEBSF before use) and sonicated (Vibra cell (75042), Bioblock Scientific) for 8 cycles, 10 seconds on, 10 seconds off at an amplitude of 34%. Lysation was cleared by centrifugation (15’000 x g for 15 min at 4 °C). 50 ul of lysate was kept for input control. A maximum of 500 ug total protein in a volume of 950 ul was loaded per 25 ul GFP-Trap agarose beads (gta, Chromotek), prior washed with ice-cold PBS (2 min at 2’500 x g) and incubated for 1 hour rotating at 4 °C. IP was washed five times with ice-cold washing buffer (1 M NaCl, 50 mM Tris, pH 7.5, 5 mM MgCl_2_, 0.01% Nonidet P-40; supplemented with 1 mM NaF, 0.5 mM DTT and 0.5 mM AEBSF before use), transferred into a new tube and washed once with ice-cold PBS. For RNA analysis, the IPed complexes were released from the beads by incubating the pellet for 10 minutes with 50ul 4% SDS in 0,1M NaHCO_3_ (same treatment was performed with the input sample). Supernatant was collected and the beads-releasing process was repeated. To reverse crosslink the formaldehyde linking, the samples were treated in 20 mg/ml Proteinase K (Roche) in PK buffer (100 mM Tris-HCl (pH 7.5), 50 mM NaCl, 10 mM EDTA) (pre-incubated for 20 minutes at 37 °C to eliminate potential RNases) at 65 °C overnight, shaking at 1000 RPM. RNA was isolated with Trizol (Invitrogen), following the manufacturer protocol.

### RT-qPCR

MiRNAs were reverse-transcribed into cDNA using miScript II (Qiagen, 218160) and RT-qPCR was performed using the miScript SYBR Green PCR Kit (Qiagen, 218073) together with the miRCURY LNA miRNA PCR Assay (Qiagen, 339306) on a StepOne Real-Time PCR System (Applied Biosystems).

### Small RNA-Seq

Small RNA Seq was performed according to the AQ-Seq protocol from ^10^ with minor modifications. In brief, for small RNA library preparation 50 to 300 ng of total RNA was ligated with randomized 3’ adapter using T4 RNA ligase 2 truncated KQ (NEB) in 1X T4 RNA ligase reaction buffer (NEB) supplemented with 20% PEG8000 (NEB). Ligated miRNA-adapter fragments were gel-purified and eluted in 0.3 mM NaCl, 2mM EDTA (pH8). After centrifugation the EtOH-precipitated fragments were resuspended in water and ligated to 0.25 µM randomized 5’ RNA adapter using T4 RNA ligase 1 (NEB) in the same buffer conditions as described above. Reverse transcription with Superscript III First-Strand synthesis supermix (ThermoFisher Scientific) and 5 µM RT Primer (RTP, TruSeq kit; Illumina) was then performed according to the manufacturer’s recommendations. After cDNA amplification the PCR amplicons representing miRNA sized inserts were gel-purified, precipitated and resuspended in water. The quality of the libraries was assessed by qPCR and Tapestation measurements and the Libraries were sequenced on Illumina sequencers (MiSeq or NextSeq 550; according to the number of pooled libraries).

### Bioinformatics analysis

miRNA-mRNA target prediction was performed using TargetFly 7.2^11^. PCA plot, Heat map and Hierarchical clustering were performed in R using DEseq2, pheatmap and stats packages, respectively. Heat map was performed on a subset of miRNA showing a DEseq2 statistical differential expression between 2 cell types and showing an expression level of more than 1000 reads per million in at least one sample. Hierarchical clustering (Hclust function using ward method) was performed on the 2D matrix of number of target sites per target gene for each neuroblast-enriched miRNA. The X-axis of the matrix was composed of miRNAs statistically enriched in neuroblasts compared to neurons and showing an expression level over 1000 reads per million in at least one neuroblast sample. The Y-axis was composed of the list of genes predicted to be targeted by at least one of the selected neuroblast miRNAs and showing an average of expression level more than 100 reads per million in neuroblast or neuron samples reported in ^12^.

### Production of DNA constructs

The prospero 3’UTR was obtained by nested PCR performed on genomic DNA from *Drosophila* larval CNS. The prospero 3’UTR [2131bp] was subcloned into the pmirGlo Dual-Luciferase expression vector (Promega) downstream of the firefly luciferase gene, to produce pmirGLO-prospero. pri-miR-1-3p (1.5kb) was amplified from genomic DNA of *Drosophila* larval CNS and was subcloned into the pCX^65^, a derivative of the pCAGGS expression vector, to produce pCX-miR-1-3p.

### Dual Luciferase Assay

HEK293 cells were cultured in DMEM containing 10% fetal calf serum (vol/vol) 24 h before transfection and transfected in triplicates with Lipofectamine 2000 reagent (Invitrogen) in OPTIMEM in a 24 well plate according to supplier’s instruction. Transfection was done in 1:1 (250ng pri-miR-1-3p: 250ng miRGLO) plasmid DNA ratio. Firefly and Renilla luciferase activities were measured 24 h after transfection with the Dual-Glo luciferase assay system (Promega) using a Luminometer (Berthold Technologies).

### Data availability

All sequencing data generated in this work is assessible under GEO No GSE240560

